# Eat or Be Eaten: Implications of potential exploitative competition between wolves and humans across predator- savvy and -naive deer populations

**DOI:** 10.1101/2022.09.29.510202

**Authors:** Ellen M. Candler, Stotra Chakrabarti, William J. Severud, Joseph K. Bump

## Abstract

1. Recolonization of predators to their former ranges is becoming increasingly prevalent. Such recolonization places predators amongst their prey once again; the latter having lived without predation (from such predators) for considerable time. This renewed coexistence creates opportunities to explore predation ecology at both fundamental and applied levels.
2. We used a paired experimental design to investigate white-tailed deer risk allocation in the Upper and Lower Peninsulas (UP and LP) in Michigan, USA. Wolves are functionally absent in the LP, while deer in the UP coexist with a re-established wolf population. We treated 15 sites each in UP and LP with wolf olfactory cues, and observed deer vigilance, activity, and visitation rates at the interface of habitat covariates using remote cameras. Such a paired design across wolf versus no-wolf areas allowed us to examine indirect predation effects while accounting for confounding parameters such as the presence of other predators and human activity.
3. While wolf urine had no effect across most metrics in both UP and LP, we observed differences in deer activity in areas with versus without wolves. Sites treated with wolf urine in the UP showed a reduction in crepuscular deer activity, compared to control/novel-scent treated sites. Further, we observed a strong positive effect of vegetation cover on deer vigilance in these sites. This indicates that simulated predator cues likely affect deer vigilance more acutely in denser habitats, which presumably facilitates predation success. Such responses were however absent among deer in the LP that are presumably naïve towards wolf predation.
4. Where human and non-human predators hunt shared prey, such as in Michigan, predators may constrain human hunting success by increasing deer vigilance. Hunters may avoid such exploitative competition by choosing hunting/bait sites located in open areas. Our results pertaining to fundamental predation ecology have strong applied implications that can promote human-predator coexistence.

## Introduction

Predators can alter prey populations directly through lethal offtake and/or behaviorally mediated indirect effects (Kuijper et al., 2013; McNay et al., 2022; Palmer et al., 2021). While some prey succumb, others avoid predation by changing their behavior or life-histories (Kohl et al., 2018; Sabal et al., 2021). Predator-effects on prey are a fundamental focus in ecology, although, dynamic predator-prey interactions in a fast-changing Anthropocene warrant more experimental examinations of the underlying mechanisms of these direct and indirect effects (Parsons et al., 2022; Scrosati, 2021). While predator populations are declining in many parts of the world (Ripple et al., 2014), some areas are witnessing their comebacks (Gigliotti et al., 2019; Jarausch et al., 2021; Scharf & Fernández, 2018). Such contrasting levels of predator presence create (and re-create) intriguing interactions. Predators recolonizing former habitats may encounter potentially naive prey that have lost anti-predatory strategies (Berger, 1999; Berger et al., 2001).

Anti-predator responses are defense strategies that prey species employ when they perceive predation risk via visual, auditory, tactile, and/or olfactory stimuli. When at risk, prey respond through multiple mechanisms either proactively, such as by increasing vigilance, avoiding risky areas, social grouping, and birth synchrony, or reactively through evasion, camouflage, and defense-herding (Flagel et al., 2016; Ims, 1990; Kohl et al., 2018; Mech, 2007; Rowland et al., 2020). Successful anti-predatory responses, while providing survival benefits, can have severe fitness costs for individuals. For example, ungulates that intensify vigilance as a response to predator cues limited durations of feeding or avoided high-risk areas even if they are rich with resources (Chamaillé-Jammes et al., 2014; Kuijper et al., 2013; Melchiors & Leslie, 1985; Sahlén et al., 2016; Wikenros et al., 2015). Furthermore, certain herbivores are known to incur major reproductive costs while courting at breeding grounds that also attract a host of predators (Cherry et al., 2016; Jhala & Isvaran, 2016). However, perception and response to predation risk have significant decision-making implications that can result from a mix of instinctive and learnt behaviors. Thus, the risk of predation can vary greatly amongst experienced prey (prey that live under constant predation pressure) versus naive prey (prey that live without or under very low predation pressure). For example, in areas where wolves (*Canis lupus*) and grizzly bears (*Ursus horribilis*) had been extirpated, moose (*Alces alces*) did not exhibit avoidance to predator olfactory or auditory cues, while those that had been exposed to both predators showed avoidance (Berger et al., 2001). Similarly, species on predator-free islands have been observed to ’adaptively forget’ anti-predatory strategies, resulting in prey naivety (Blumstein et al., 2004)

The Upper and Lower Peninsulas (UP and LP) of Michigan, USA offer a novel opportunity to explore these drivers of predator-effects. In the LP, white-tailed deer (*Odocoileus virginianus*) occur where wolves have been absent since the mid-1800s. Individual wolves have been detected (although rarely) in the LP but remain functionally absent as predators of deer. Alternatively, deer in the nearby UP coexist with a re-established wolf population since the mid-2010s (O’Neil et al., 2017). Both peninsulas have a similar guild of other deer predators – black bears *U. americanus*, coyotes *C. latrans*. Also, white-tailed deer are a subsistence species for indigenous communities and a highly valued hunting species in this region. In both UP and LP, deer are typically hunted by humans at specific locations where deer are food conditioned with bait consisting of corn, fruits, or vegetables – creating artificially provisioned resource rich sites on the landscape. These resource-rich but localized hunter-bait sites are intended to attract and condition deer to revisit these sites frequently. According to behavioral optimality, prey species need to balance the conflicting demands of efficient foraging and predator avoidance (Lima & Bednekoff, 1999; Sih, 1980). Thus, deer behavior at these hunter-bait sites is expected to be mediated by the perceived risk of predation because wolves are attracted to bait sites as well (Bump et al., 2013), which in turn should be influenced by prey experience or naivety towards predators.

Through a paired experimental design, that accounted for confounding effects of other predators and human hunters, we tested for differences in risk perception by examining the immediate behavioral effects of wolf olfactory cues on naive versus experienced deer at hunter-bait sites. Since we expected no response from naive deer (LP deer; no functional wolf population) to wolf scent (urine), we predicted:

1. deer that generally experience wolf predation (UP) would increase group size, vigilance intensity, and occur less often at bait sites with wolf cues (scent),
2. the immediate effects of predator cues on deer vigilance will be further modulated by habitat covariates such as vegetation cover at the sites, with higher vigilance to be expected in denser habitats. Prey are often cautious, and tend to avoid ‘risky’ areas that lack good visibility and offer ambushing opportunities for predators even where food is abundant (Edwards, 1983; Kuijper et al., 2013; Ripple & Beschta, 2004),
3. predation-experienced deer would adjust their activity patterns to avoid crepuscular times when wolves are most active, while naive deer would not make such adjustments. This prediction is in accordance with the predation risk allocation hypothesis that suggests temporal variation in predation risk severity affects foraging and vigilance behavior in prey (Lima & Bednekoff, 1999).

## Materials and Methods

### Study Area

We conducted this study in Michigan’s UP and LP from September 21, 2018 to November 10, 2018. The UP study area was entirely within the Hiawatha National Forest and was interspersed with unpaved, US Forest Service roads. The bait sites in the LP study area were all located in Mackinaw State Forest of Michigan, and the area was also interspersed with private agriculture, forestland, and paved, secondary roads (Fig. 1). The main vegetation cover, derived from the National Land Cover Database, at the UP study sites was evergreen (5 sites), woody wetland (3 sites), grassland (3 sites), deciduous (2 sites), and mixed forest (2 sites), while the LP study sites were deciduous (6 sites), scrub/shrub (4 sites), grassland (3 sites), and evergreen (2 sites; Fig. 1) (Homer et al., 2015).

**Figure 1.**
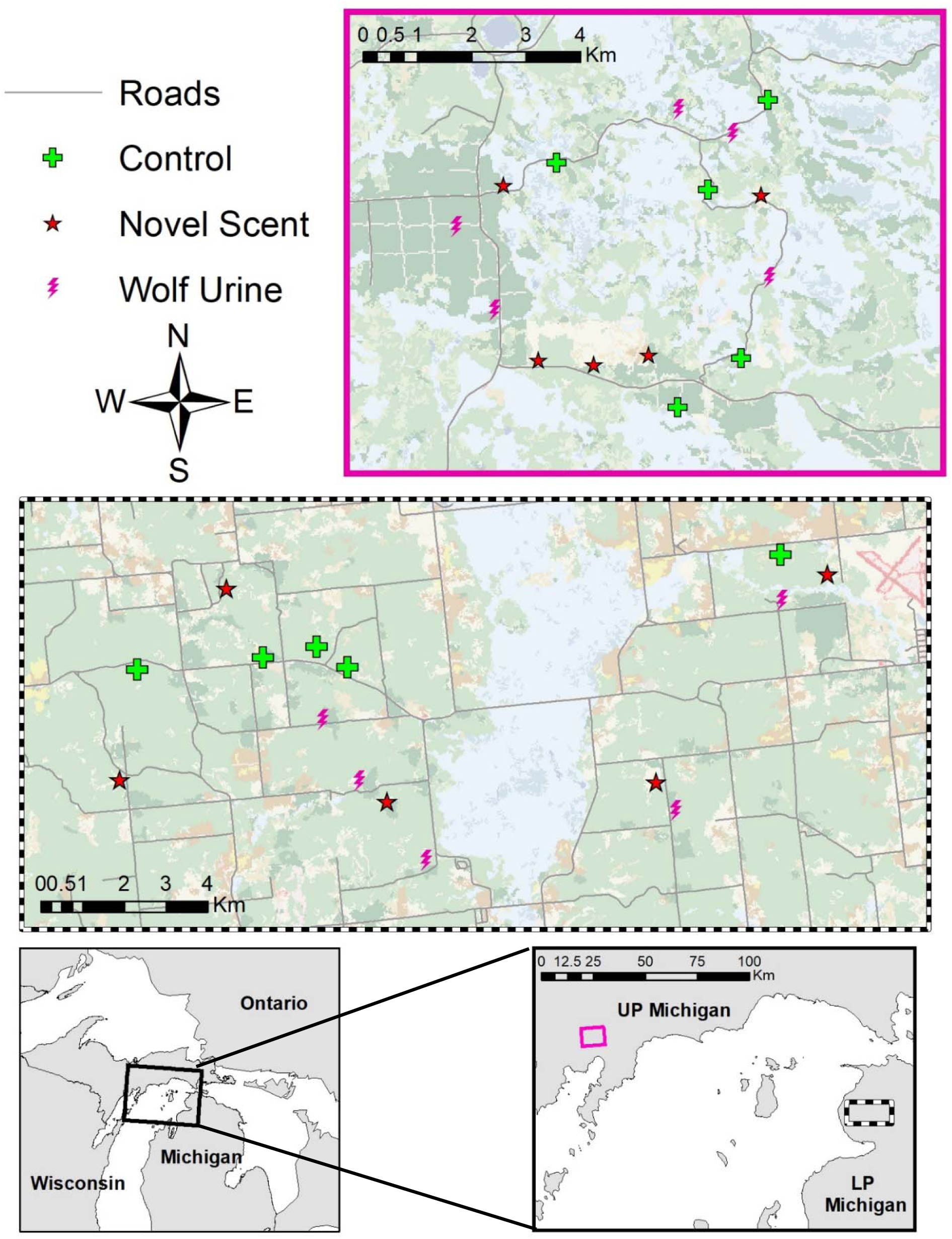
Locations of 30 experimental deer bait sites with camera traps deployed between September and November 2018 in Michigan, USA. The top inset map (solid magenta line) indicates the 15 sites in the Upper Peninsula (UP; wolves [*Canis lupus*] present) and the lower inset map (black dashed line) indicates the 15 sites in the Lower Peninsula (LP; wolves absent). The different symbols indicate the different treatments (wolf urine, lemon juice and distilled water) assigned to each site.

### Study Design

We established 30 deer bait sites between the two study areas, 15 in the UP and 15 in the LP (Fig. 1). To mimic recreational hunters, we selected sites based on deer hunting desirability such as habitat openness and/or proximity to a deer trail (Peterson, 2015). Sites were at least 1 km away from each other to avoid duplication and dependence (Winke, 2012).

We baited each site with 3.8 liters of corn spread in a 3 by 3-meter-square plot, as per the Michigan deer baiting regulations (Michigan Department of Natural Resources, 2018). For the duration of the study, all sites were baited each week in order to maintain 3.8 liters of corn on the ground to replicate actual hunter behavior and to reduce the confounding effect of bait availability on deer behavior. We deployed one remote camera (Reconyx Hyperfire series or Cuddeback E) on a tree 4 meters from the center of the bait site. Each camera was programmed to take 5 burst images with no delay between triggers to record the number of deer present, their posture, sex (when possible), and presence/absence of fawn(s). We replaced batteries and memory cards weekly.

#### BACI Experimental Setup

We used a 10 week before-after-control-impact (BACI) design for the experimental treatment of both areas. This BACI design was conducted so that all sites were paired with themselves with all other factors (e.g., other predators, human visitation, environmental effects) remaining comparable. This allowed us to tease apart the effects of the treatment. Before experimental treatments, we collected data for one week preceding the six-week period to establish a baseline condition of deer visitation to the bait sites (Stewart-Oaten et al., 1986). For the first 3 weeks (September 28-October 20), we baited sites and recorded deer occurrence and behavior via remote cameras with no scent treatments applied. For the following three weeks we continued to bait and record deer after treating the sites with three scents. In each area, UP and LP, 5 sites were treated each with wolf urine (predator treatment), lemon juice (novel scent; not a native plant), or distilled water (control). We assigned site treatment randomly (Atkins et al., 2016; Wikenros et al., 2015). When we treated sites in the field, we visited all ‘control’ sites in an area first followed by the ‘novel scent’ sites, and finally ‘experimental/wolf urine’ sites to avoid cross-site spread of scents. We also used different treatment application tools (i.e., pipets, travel containers) for each treatment type to avoid cross contamination of treatments. At each site, we applied 10 mL of the given treatment (experimental, novel, or control) by dripping it from a pipet onto the bait site. This was done to mimic the amount and application of scent marking of a pair of wolves (Peters & Mech, 1975).

#### Vegetation Cover

We also recorded predator concealment or hiding potential (horizontal cover) at each site to account for site-specific differences in habitat openness - an index of perceived predation threat. We used a 2 m cover pole with 20 sections, each 10 cm long, to measure predator hiding potential at each site (Kuijper et al., 2014; Severud et al., 2019). From the center of each bait pile, we measured the horizontal cover in each cardinal direction. While one person held the cover pole 10-meters away from the center of the bait pile, another took a picture of the cover pole with a camera stationed on a 1-meter high pole at the center of the bait pile, mimicking deer visual height (Severud et al., 2019). We conducted this procedure twice during the study, once at the beginning and once when treatment started, to account for change in hiding potential with loss of leaves later in the year. For each of the 20 (10 cm) sections on the 2 m cover pole, we estimated obscurement to the nearest 25% (Severud et al., 2019). We calculated a single mean and standard error value for each site and each measurement time (beginning and middle of study).

### Analysis

#### Photo Analysis

Images were tagged in batches using the DigiKam photo editing software (Niedballa et al., 2016). Each batch was defined by any set of images taken within 5 minutes of the prior image. In each batch, for deer images only, we recorded if fawns were present, labelled all sexes when possible, and tallied a total count. This increased the likelihood that we accounted for all individuals that visited the bait pile and avoided miscounting any that were out of the camera frame. For individual pictures within batches, in addition to generic behaviours (e.g., fighting, nursing, foraging), we recorded the number of individuals with their heads above their shoulders, indicating vigilance/state of alertness (Fig. 2; Flagel et al., 2016; Schuttler et al., 2017). We also labelled batches for different species that were captured at site visits including squirrels (*Sciurus* spp.), raccoons (*Procyon lotor*), wild turkeys (*Meleagris gallopavo*), and coyotes. We did not detect additional scent marking by other species.

**Figure 2.**
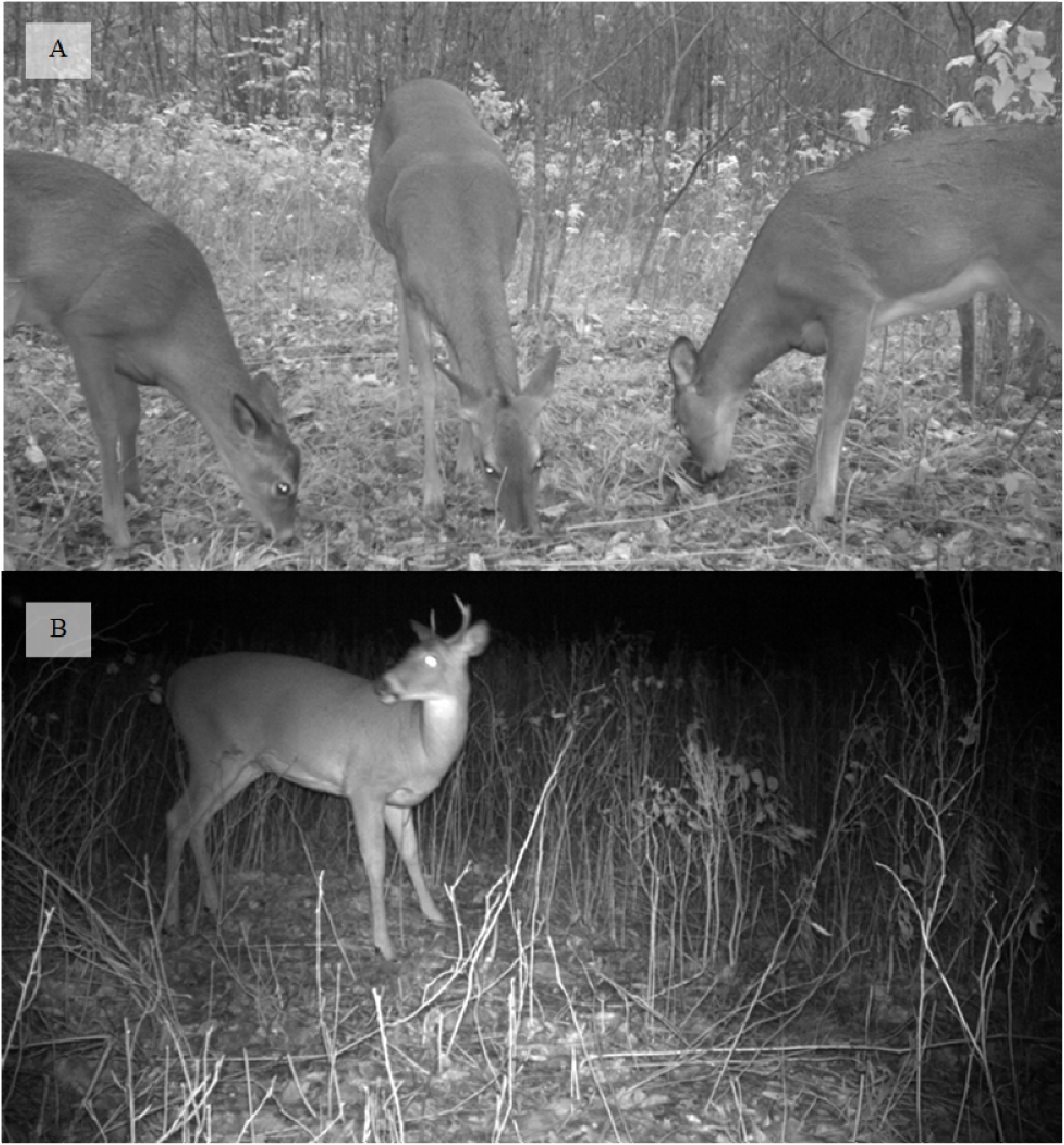
Camera-trap detections of white-tailed deer (*Odocoileus virginianus*) at simulated deer hunter-bait sites September to November 2018 in Michigan, USA. The representative images indicate vigilance (A) and non-vigilance (B) behavior.

#### Treatment Impact on Deer Vigilance, Group Size and Visitation Rate

We explored the impact of different treatments in the two areas (UP and LP) on the number of deer at each site, proportion of group that was vigilant, and the number of visits made to the site. To account for the paired nature of our treatment design, we first averaged the variable values for each site individually and calculated a difference in values before and during treatment application. We then averaged across all sites within a given treatment. We also calculated a vigilance intensity metric using equation 1. This metric includes both group vigilance and event duration.

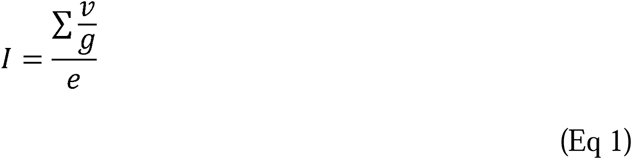

Where *I* is the vigilance intensity metric for a given event, *v* is the number of deer vigilant in a single image, *g* is the group size in a single image, and *e* is the total time of the event. Hence, vigilance intensity simply standardizes individual vigilance across different group sizes and time spent in front of the camera. Similar to the previous analysis, we calculated a difference in vigilance intensity before and after treatment for each treatment type in each peninsula.

#### Diel Activity

To analyze possible temporal variability in the use of bait sites in the UP and LP before and after treatment, we used nonparametric kernel density estimation (Prugh et al., 2019; Wang et al., 2015). We converted times to radians and used a kernel density estimator to create a probability density distribution for each before or after period (Ridout & Linkie, 2009). We calculated the proportion of temporal overlap between the two treatment periods for each treatment in an area (Wang et al., 2015). We used a Δ◻_4_ with a smoothing parameter of 1 because our sample size for all analyses was greater than 50 (Ridout & Linkie, 2009). We conducted this analysis using the overlap package (Meredith & Ridout, 2018; Wang et al., 2015) in R (R Development Core Team 2013).

We applied Watson’s U2 statistic with the CircStats package to test for homogeneity between the two samples of interest (i.e., test for/detect a statistically significant shift in the diel pattern before and after treatment) (Lashley et al., 2018; Lund & Agostinelli, 2012). If deer significantly shifted their temporal pattern between the two treatment periods, we expected Watson’s U2 statistic would be greater than the critical value (0.19 for an α value of 0.05) and P < 0.05. In the UP, we predicted that there would be no shift in temporal visitation by deer at the control and lemon treated sites (high Δ◻_4_, U2 ≤ 0.19), while a shift to more nocturnal activity at the wolf urine treated sites would be detected (lower Δ◻_4_, U2 > 0.19; Kohl et al. 2018) to potentially avoid wolves that are known to be typically less active at night (Kohl et al., 2018). In the LP, we expected that we would not see a significant shift in any treatment sites because deer in the LP are ostensibly naive to wolf predation.

#### Effect on Vigilance Behaviour

We fitted generalized linear models using pooled data from both the UP and LP to check for the effect on overall vigilance intensity by treatment effect (before/after) and type (predator/novel/control), area (UP or LP) and vegetation cover (i.e., predator hiding potential; S1; Fležar et al., 2019; Prugh et al., 2019) through additive and interactive effects between parameters of interest. Subsequently, we fitted models to examine the relationship between vigilance intensity and vegetation cover at each treatment type within both the UP and LP before and after treatment.

Using generalized linear model, we further examined the effects of sex, vegetation cover, and presence of young on vigilance intensity, and included additive and interactive effects with area to detect any difference between the UP and LP. We ranked models based on AICc.

### Results

We obtained 286,436 images over seven weeks. After removing images taken during the pre-experimental baseline week and images that did not contain deer, we retained 213,264 images for analysis with 85,675 images taken in the LP and 127,589 images taken in the UP.

#### Impact on Deer Vigilance, Group Size and Visitation Rate

We observed a near 30% increase in group size at sites treated with wolf urine in the UP but this increase translates to less than half a deer - an increase that is not ecologically meaningful. Additionally, we observed a slight increase (<3%) in the number of visits at both lemon and control treated sites in the UP and a marginal decrease (<4%) in the proportion of the group that was vigilant at lemon treated sites in the LP (Table 1, Fig. 3). We also observed a decrease in vigilance intensity of deer at wolf urine treated sites in the LP (Fig. 3). We did not detect a significant change before and after treatments for any other treatment sites in either area for the four response variables, i.e., the proportion of group that is vigilant, number of visits/visitation rate, group size and vigilance intensity (Fig. 3).

**Figure 3.**
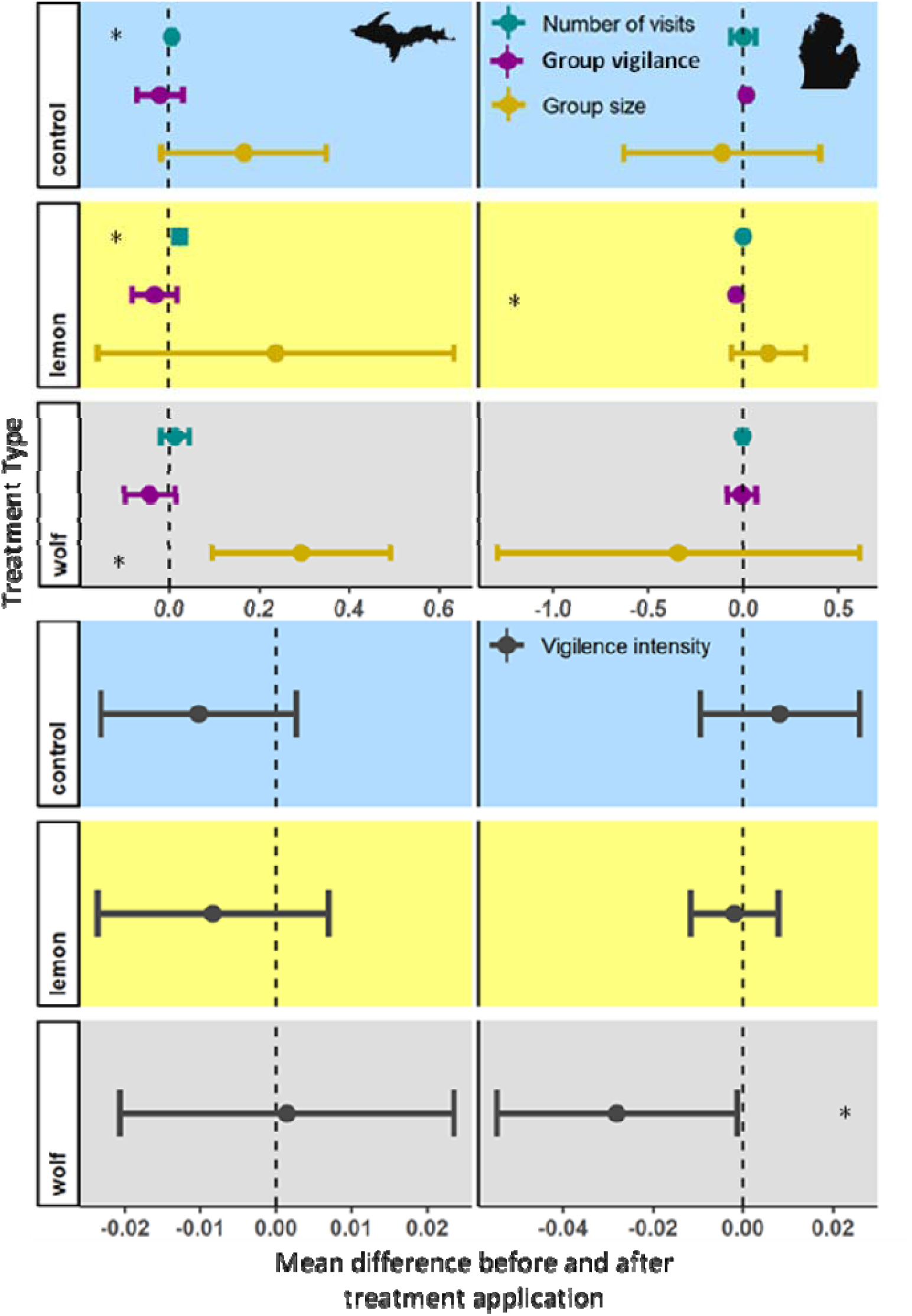
Perceived effect of wolf (*Canis lupus*) predation risk on deer behavior. Difference before treatment period and during treatment period in group size, group vigilance, number of visits, and vigilance intensity (sum of proportion of group vigilance in each event divided by total time of event) from September to November 2018 in Michigan, USA. Symbols indicate the mean difference and error bars indicate 95% confidence interval. Blue shading represents the control sites (water), yellow indicates sites treated with novel scent (lemon), and grey indicates wolf-urine treated sites. Positive values indicate an increase in the metric. Asterix indicates p < 0.05.

**Table 1.** Perceived effect of wolf (*Canis lupus*) predation risk on deer behavior. Mean difference before treatment period and during treatment period in group vigilance, group size, number of visits, and vigilance intensity (sum of proportion of group vigilance in each event divided by total time of event) from September to November 2018 in the Upper Peninsula (UP) and Lower Peninsula (LP) Michigan, USA. *Differences before treatment period and during treatment periods were significant if the 95% confidence interval did not include 0.

| Area | Response variable | Treatment | Mean difference | 95% CI |
| --- | --- | --- | --- | --- |
| UP | Group vigilance | wolf | -0.044 | -0.101 to 0.013 |
|  |  | lemon | -0.032 | -0.082 to 0.017 |
|  |  | control | -0.020 | -0.071 to 0.031 |
|  | Group size | wolf | 0.293* | 0.093 to 0.492 |
|  |  | lemon | 0.238 | -0.159 to 0.635 |
|  |  | control | 0.166 | -0.018 to 0.351 |
|  | Number of visits | wolf | 0.011 | -0.020 to 0.043 |
|  |  | lemon | 0.024* | 0.011 to 0.037 |
|  |  | control | 0.007* | 0.003 to 0.011 |
|  | Vigilance intensity | wolf | 0.001 | -0.021 to 0.024 |
|  |  | lemon | -0.008 | -0.024 to 0.007 |
|  |  | control | -0.010 | -0.023 to 0.003 |
| LP | Group vigilance | wolf | -0.007 | -0.083 to 0.068 |
|  |  | lemon | -0.035* | -0.044 to -0.025 |
|  |  | control | 0.016* | 0.002 to 0.034 |
|  | Group size | wolf | -0.341 | -1.294 to 0.611 |
|  |  | lemon | 0.133 | -0.061 to 0.327 |
|  |  | control | -0.110 | -0.626 to 0.405 |
|  | Number of visits | wolf | 0.004 | -0.019 to 0.012 |
|  |  | lemon | 0.001 | 0.002 to 0.004 |
|  |  | control | 0.000 | -0.065 to 0.065 |
|  | Vigilance intensity | wolf | -0.028 | -0.055 to 0.001 |
|  |  | lemon | 0.002 | -0.012 to 0.008 |
|  |  | control | 0.008 | -0.010 to 0.026 |

#### Diel Activity

Deer changed their diel activity in the LP and UP, and in all treatment types, before and after treatment application (i.e., all U2 > 0.19). We did not detect a consistent directional pattern in the temporal shifts for most sites. However, we did observe a difference between the UP and LP diel patterns wherein the deer in the UP tended to reduce activity between 6:00 and 18:00 both before and after treatment application while the deer in the LP did not have a consistent inactivity period (Fig. 4). Additionally, deer at wolf-treated sites in the UP reduced bait site visitation in the evening hours (i.e., 1800h; Fig. 4).

**Figure 4.**
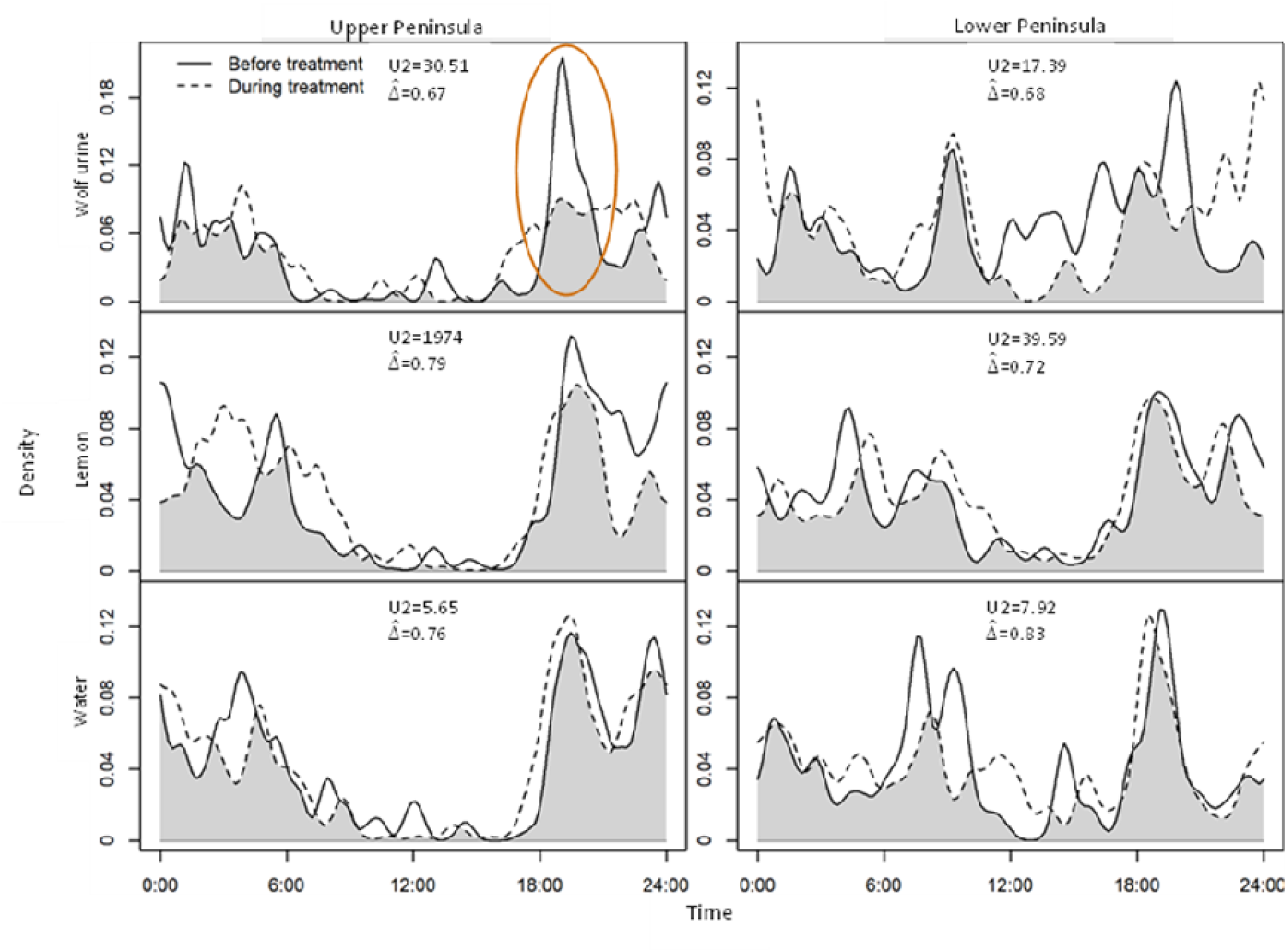
Temporal activity patterns of white-tailed deer (*Odocoileus virginianus*) populations before (solid line) and during treatment application (dashed line) in the Upper Peninsula (UP) and Lower Peninsula (LP) of Michigan, USA between September and November 2018. Shaded area indicates the temporal overlap between the 2 periods. The orange circle highlights the decrease in deer activity at wolf (*Canis lupus*) urine treated sites in the UP. The reported U2 statistic is evaluated against the test statistic U2 = 0.19.

#### Impact of Vegetation Cover

Deer vigilance intensity increased by 1.3 times for each unit increase in vegetation cover in the UP but not in the LP (Fig. 5; S1). Though vigilance intensity did not increase with vegetation cover in the LP, the base vigilance intensity (model intercepts) increased significantly after all treatment applications (S2).

**Figure 5.**
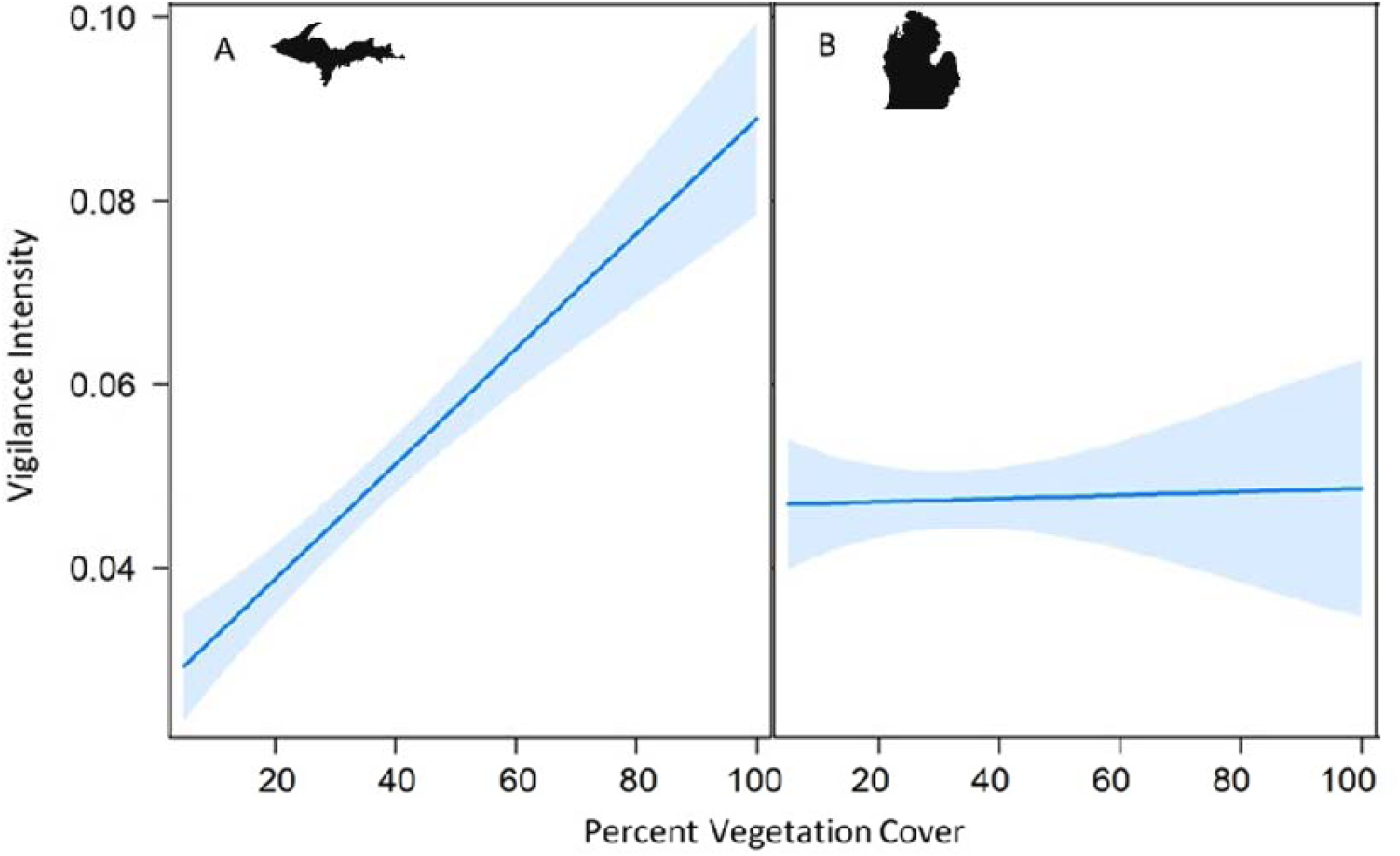
Effects graph showing the predicted vigilance intensity (sum of proportion of group vigilant in each event divided by total time of event) of white-tailed deer (*Odocoileus virginianus*) by percent vegetation cover for all treatment sites across the Upper Peninsula (UP) of Michigan, USA (A; wolves present); and all treatment sites across the Lower Peninsula (LP) of Michigan, USA (B; wolves [*Canis lupus*] absent).

We detected an increase in vigilance based on sex (males showing higher vigilance than females) and when young were present, however, these effects did not vary between the UP and the LP (Table 2). The most parsimonious model from our analysis indicates that the variation in vigilance intensity was caused by difference in vegetation cover, modulated by area (Table 2, model 1).

**Table 2.**
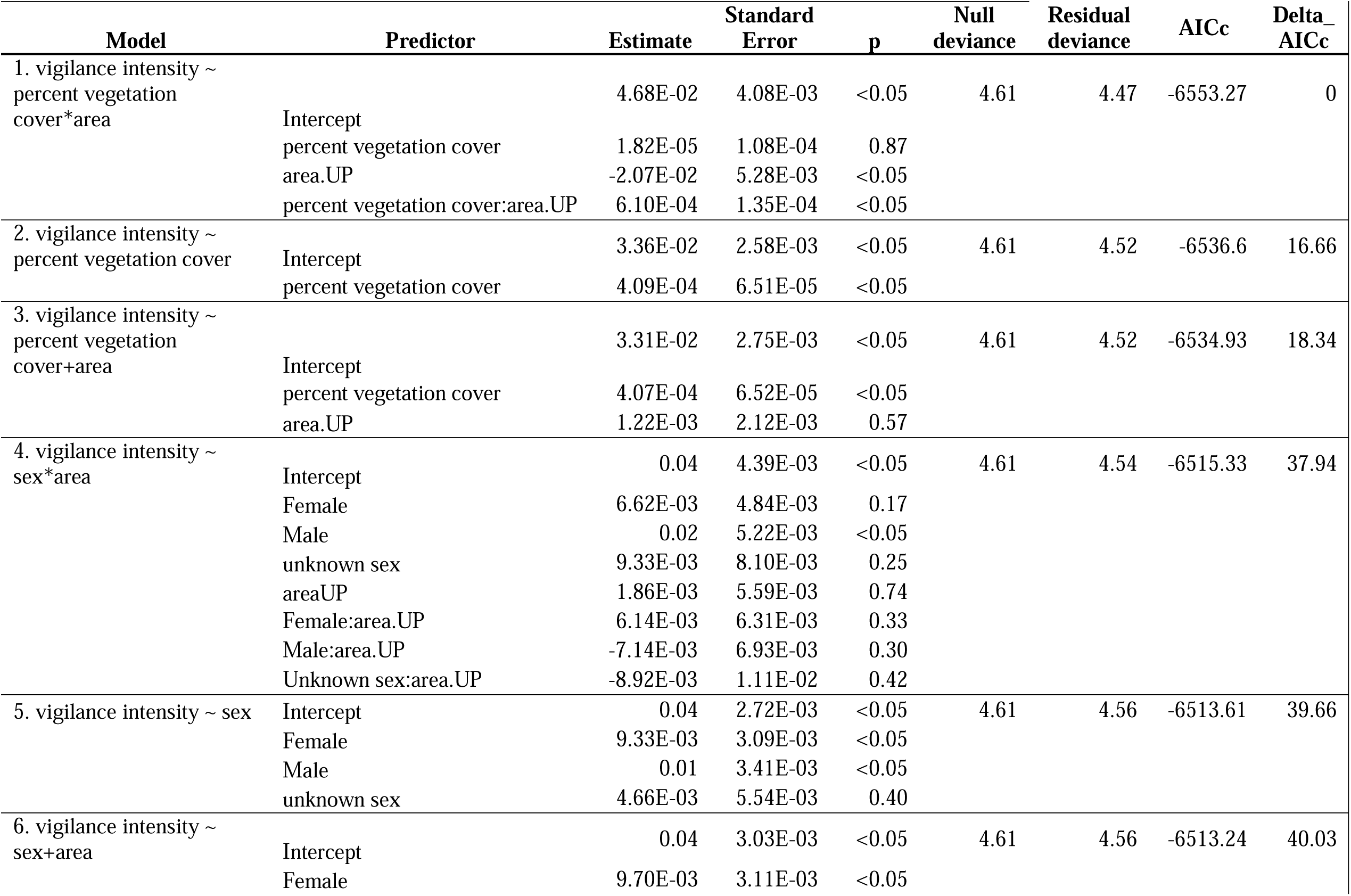
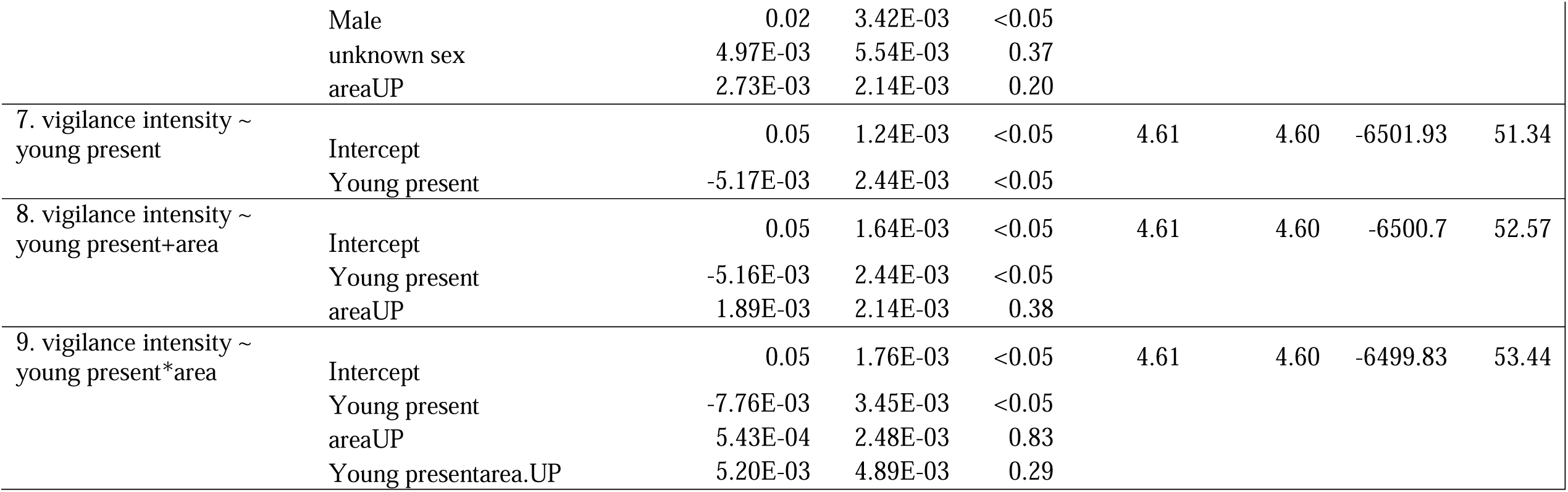
Performance of nine individual, additive, and interactive models predicting vigilance intensity for white-tailed deer (*Odocoileus virginianus*) from September to November 2018 in the Upper Peninsula (UP) and Lower Peninsula (LP) Michigan, USA.

## Discussion

Predator absence for long durations can lead to the gradual erosion of anti-predatory responses of prey (Berger, 1999; Berger et al., 2001). Subsequent recolonization of predators to such landscapes raises questions about the response that naive prey will have once predators return, particularly in prey populations valued and pursued by human hunters. We investigated the effects of simulated predator presence on experienced versus naive deer at food-rich hunter-bait sites in Michigan. We did not find a significant effect of wolf scent treatment on deer vigilance, number of visits, or vigilance intensity among predator-savvy/experienced deer in the UP but did note a slight increase in group size, though the difference was not ecologically meaningful. A similar lack of response to wolf urine has been documented in deer populations in the Netherlands and Poland where little support was found for effects of predator scents in both predator savvy and predator naive prey populations even when three times the volume of wolf urine was used (van Ginkel, Smit, et al., 2019). This lack of reaction to wolf scent cues in both areas may also be a result of decreasing response over time to the wolf urine as it degrades (Bytheway et al., 2013; Kuijper et al., 2013; Wikenros et al., 2015). Introducing cues that indicate immediate wolf proximity, such as howling or visual, are likely to produce different results (Liley & Creel, 2008; van Ginkel, Smit, et al., 2019). The absence of any response to wolf urine may also be a result of predator hunting strategies. Cursorial predators might fail to evoke strong anti-predatory responses in deer, as opposed to pronounced effects as observed in the case of cues from ambush predators (Preisser et al., 2007; Sunde et al., 2022).

We also observed a significant shift in diel patterns at all sites before and after treatment but there was no consistency to the shift in activity for most treatment sites. However, deer at sites treated with wolf scent in the UP (where they are sympatric with wolves) reduced their visitation in the evening hours. This finding is in accordance with other research that suggests that ungulate prey shift their activity to avoid riskier times (Gehr et al., 2018; Kohl et al., 2018). For example, deer in Minnesota, USA demonstrated a more diurnal pattern under the effects of wolf scent to likely avoid the riskier crepuscular times when wolves tend to hunt (Palmer et al., 2021). It is also noteworthy that before and after treatment application, all UP sites exhibited a decrease in deer activity at bait sites between 6:00 and 18:00 hours, while the LP sites had inconsistent patterns. This suggests that deer in the UP are perhaps avoiding peak wolf activity/hunting hours regardless of predator cues at sites, while LP deer do not have as large of an inactivity period, likely because they do not coexist with wolves (S2; Kohl et al., 2018).

Our results mirror findings from similar studies that found that deer vigilance was dependent on sex and fawn presence (Cherry et al., 2015; Gulsby et al., 2018; Higdon et al., 2019; Lashley et al., 2014). When young were present, vigilance increased regardless of wolf presence on the landscape. Similarly, sex played an important role where males showed more vigilance than females (Table 2). However, we found that vegetation cover was the largest contributor to the variance in vigilance, and this difference in vigilance was modulated by wolf presence or absence in that area (Cherry et al., 2015; Olson et al., 2021).

Our results show a strong effect of vegetation cover on deer vigilance intensity where wolves are established. Increase in vegetation cover at the UP sites were characterized by intense vigilance activity in deer, regardless of the treatment. Denser vegetation likely represents more predator hiding potential and difficult escape routes, likely causing an increase in deer vigilance regardless of olfactory cues at the site (Fig. 5; Dellinger et al., 2019; van Ginkel, Kuijper, et al., 2019; Wikenros et al., 2015). Research comparing white-tailed deer and mule deer (*Odocoileus hemionus*) in Oregon’s expanding wolf range reported that white-tailed deer did not typically avoid areas with wolves but did favor areas with less dense vegetation that facilitates an easy escape because they relied on flight and early detection of predators (Dellinger et al., 2019). Other studies also demonstrated the importance of escape routes for white-tailed deer, indicating that white-tailed deer flee rather than stand their ground in the presence of a predator (Lingle & Pellis, 2002). This phenomenon has also been noted in other species such as chacma baboons (*Papio ursinus*) in Gorongosa National Park, Mozambique where vocalization and vigilance were heightened in denser habitats (Hammond et al., 2022). Our findings further highlight the importance of considering prey behavior and escape strategies, along with habitat characteristics for a mechanistic understanding of predator-effects.

While other studies, investigating questions similar to ours, have found that sex and presence of young were the strongest predictors of deer vigilance, several key aspects in our research differed from these studies, and might explain our subtly different results (Cherry et al., 2015, 2017; Gulsby et al., 2018; Higdon et al., 2019; Lashley et al., 2014). In the cited studies, vegetation was standardized or the time of year the research was conducted was different than our study period. If we were to conduct this experiment again during spring when fawns are more susceptible to coyote predation, the influence of vegetation may be masked with intensified vigilance because coyotes occur in both areas. Consequently, a season-specific antipredator response, recorded in many of the aforementioned studies, could potentially be present in our study system (Cherry et al., 2017; Gulsby et al., 2018).

Though vegetation cover did not have a significant effect at LP sites, deer at sites treated with wolf urine demonstrated higher baseline vigilance intensity after treatment was applied but intensity did not vary across vegetation cover. This likely suggests that naive deer may demonstrate an innate fear of wolves, also reported in East Limestone Island, British Columbia, Canada (Chamaillé-Jammes et al., 2014). However, when the novel scent (lemon) and control sites were assessed in the same context, we observed a similar pattern indicating that something other than predator scent might be affecting baseline vigilance later in the study. Because the second half of the study was conducted during archery hunting season, this shift in vigilance may indicate a response to increased human presence (Benhaiem et al., 2008). A study in Poland found that red deer (*Cervus elaphus*) demonstrated heightened vigilance during hunting season when hunters were likely pursuing them (Proudman et al., 2020). Hunting seasons occurred simultaneously in the UP and LP making our study an appropriate comparison between the two locations. Similarly, other predators (e.g., black bears) occur in both areas, reducing any confounding effects such predators might have had. This shift in baseline vigilance could also be attributed to the onset of the deer breeding season, an observation made in other white-tailed deer studies (Cherry et al., 2015; Schuttler et al., 2016).

Our findings offer partial support to the predator risk allocation hypothesis by demonstrating a shift in activity by prey with predator experience to a less risky feeding time - when predators are less active (Kohl et al., 2018; Lima & Bednekoff, 1999). This hypothesis may also explain the lack of difference in vigilance detected in the UP-deer population in relation to the wolf urine treatment. Because this population is under constant predation pressure from wolves and frequently encounters wolf cues, deer in the UP may demonstrate a change of vigilance especially at food rich locations such as hunter-bait sites, in order to take advantage of the food source as quickly as possible. Deer in the UP are also likely, as we observed, to show more vigilance in areas with higher predation potential, such as with dense vegetation cover (Dellinger et al., 2019; Kunkel & Pletscher, 2001).

Finally, through our experimental study we highlight the importance of considering both spatial and temporal factors when understanding predator effects on experienced versus naive prey. Past research has suggested that naive prey do not maintain the innate ability to react to predator olfactory cues but do have the ability to learn and adjust to predators within one generation (Berger et al., 2001; Steindler et al., 2020). Though predator olfactory cues did not have a significant effect on most variables we investigated, we observed a strong predator effect depending on vegetation cover that ostensibly modifies risk.

The use of a BACI experimental method in a predator free and predator savvy population allowed us to account for the complexity of the systems and make a few key conclusions: 1. Wolf scent had little effect on number of visits, group vigilance, or group size but it did impact the time that predator-savvy deer visited bait sites (deer decreased visitation in the evenings), 2. The overall diel patterns of predator-savvy deer generally avoided bait visitation between 6:00 and 18:00, whereas predator naïve deer did not show a similar avoidance pattern, and 3. As predator hiding potential (vegetation cover) increased, so did vigilance intensity among predator-savvy deer, but not in the predator-naïve deer. Because the only predator difference between the two populations is wolves, we can suppose this difference in vigilance intensity is due to wolf presence.

The recreational hunting tradition of baiting is controversial inside and outside the hunting community. Baiting increases the opportunity for a successful shot and more immediate kill, but also creates a predictable food source that concentrates deer and deer scent, which in turn attracts wolves (Hughes et al., 2010). As large predators continue to recolonize their former ranges, they will inevitably overlap with human hunters (Bled et al., 2022; Grima et al., 2021). Our current research can shed light on the impact a predator has on a predator savvy and predator naive prey population as it relates to hunting by humans. We show that predator cues can affect deer vigilance at bait-sites with denser vegetation, seemingly making deer cautious and difficult to hunt, thereby potentially affecting the experience and satisfaction reported in hunter surveys. However, we strongly urge readers, managers, and the hunting community to interpret our results cautiously and constructively. Our results can be formative in aiding carnivore-human coexistence wherein human hunters can have more success if they potentially choose not to bait/hunt deer at vegetatively dense sites.

## Author Contributions

Ellen M. Candler conceived the idea, developed the sampling design and methodology, preformed data collection, conducted the formal analysis, oversaw project administration, acquired resources, supervised field work, validated, and visualized data, and wrote, reviewed, and edited the original draft. Stotra Chakrabarti helped develop the sampling design, and methodology, analyzed, validated, and visualized data, wrote, reviewed and edited drafts; William J. Severud developed the sampling design and methodology, reviewed and edited drafts, Joseph K. Bump conceived the idea, developed the sampling design, was in charge of project administration and supervision, reviewed and edited drafts, acquired funding and resources.

## Statement on inclusion

The team of authors includes members representing both the Global North and South countries, including scientists based in the country where the study was carried out. Furthermore, both the lead and the last authors have significant association with the state where this study was conducted. All authors were engaged early on with the research and study design to ensure that the diverse sets of perspectives they represent was considered from the onset. Whenever relevant, literature published by scientists from the region as well as from other parts of the world were cited.

## Acknowledgements

This material is based upon work supported by the National Science Foundation Graduate Research Fellowship Program under Grant No. 00039218. Any opinions, findings, and conclusions or recommendations expressed in this material are those of the authors and do not necessarily reflect the views of the National Science Foundation. This research was also supported by grants from the National Science Foundation to J. K. Bump (NSF ID#1545611, NSF ID#1556676). We thank M. Dean, K. Rajski, and A. Schubel for field assistance.

## Data Availability Statement

Data has been uploaded to the Dryad Digital Repository. https://datadryad.org/stash/share/LhDJMgwhDbAX3LDhATW7XPgJHpHdfZQRIAIB5ITfwQo

