## Supplemental table and figure for "Eat or Be Eaten: Implications of potential exploitative competition between wolves and humans across predator- savvy and -naive deer populations"

S1. GLM models and outputs that predict vigilance intensity for deer September to November 2018 in Michigan, USA.

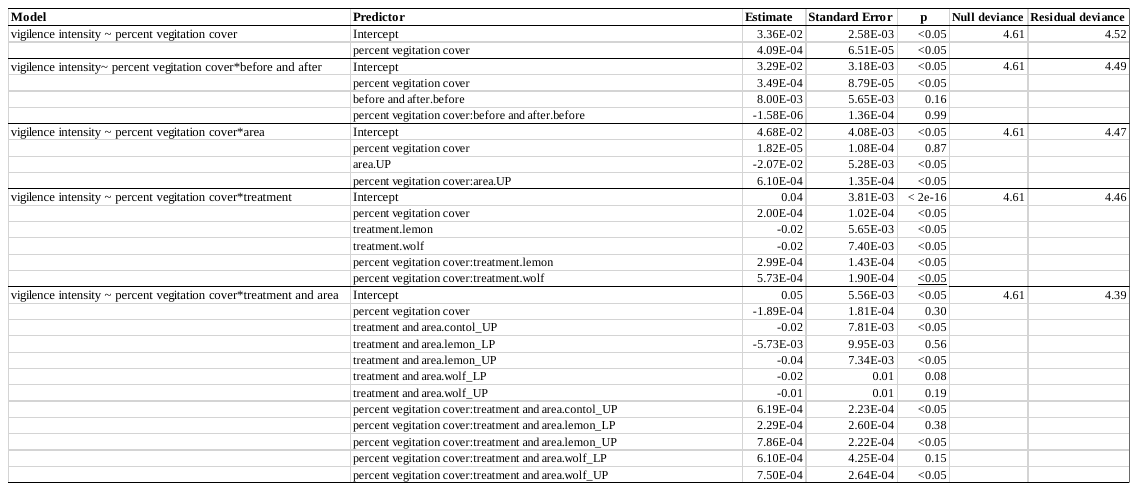

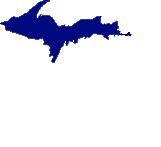

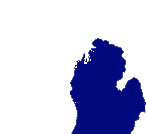

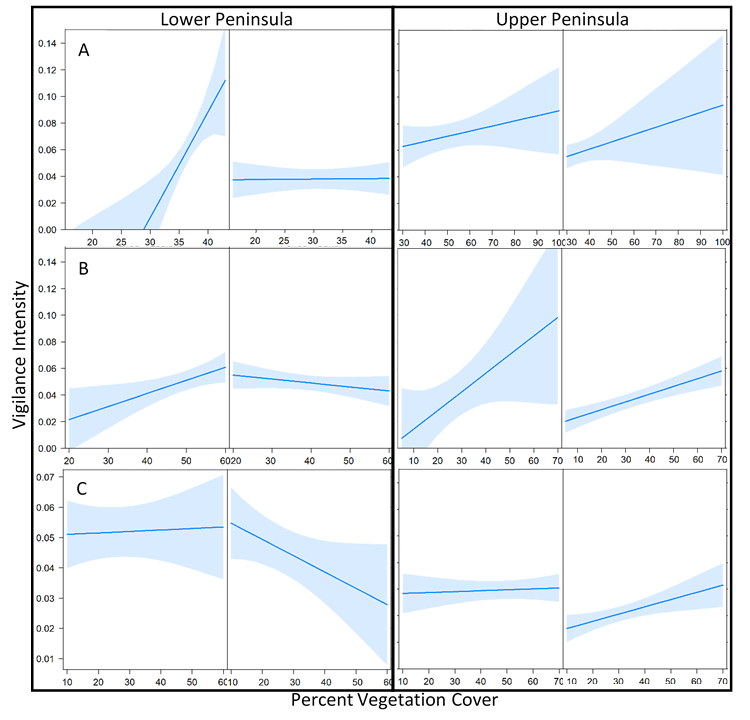

S2. Effects graph showing the predicted vigilance intensity (sum of proportion of group vigilant in each event divided by total time of event) of white-tailed deer by percent vegetation cover for wolf urine treatment sites (A); lemon treatment sites (B); and control treatment sites across the Upper Peninsula Lower Peninsula of Michigan, USA (C).
